# N6-methyladenine DNA demethylase ALKBH1 regulates mammalian axon regeneration

**DOI:** 10.1101/2020.08.19.258038

**Authors:** Qiao Li, Cheng Qian, Harry Feng, Tyger Lin, Ying Huang, Feng-Quan Zhou

## Abstract

Recent studies have shown that DNA N6-methyladenine (N6-mA) modification is emerging to be a novel and important epigenetic regulator of mammalian gene transcription. Several studies demonstrated DNA N6-mA in human or rodents was regulated by methyltransferase N6AMT1 and demethylase ALKBH1. Moreover, studies in mouse brain or human glioblastoma cells showed that reduced level of N6-mA or higher level of ALKBH1 was correlated with up regulated levels of genes associated with neuronal development. We thus investigated the functional roles of ALKBH1 in sensory axon regeneration. Our results showed that ALKBH1 regulated the level of N6-mA in sensory neurons, and upon peripheral nerve injury ALKBH1 was up regulated in mouse sensory neurons. Functionally, knocking down ALKBH1 in sensory neurons resulted in reduced axon regeneration *in vitro* and *in vivo*, which could be rescued by simultaneously knocking down N6AMT1. Moreover, knocking down ALKBH1 led to decreased levels of many neurodevelopment regulatory genes, including neuritin that is well known to enhance axon growth and regeneration. Our study not only revealed a novel physiological function of DNA N6-mA, but also identified a new epigenetic mechanism regulating mammalian axon regeneration.

**Significance Statement:** The study demonstrated that DNA N6-methyladenine (N6-mA) modification played important roles in regulation of sensory axon regeneration, likely through controlling the expression of neurodevelopmental associated genes. The results will add new evidence about the physiological function of DNA N6-mA and its regulatory demethylase ALKBH1 in neurons.

## Introduction

DNA methylation is a key epigenetic regulatory approach for many biological processes, such as genomic imprinting, epigenetic memory maintenance, aging, and neural development. The most well-known DNA methylation in mammals is 5-methylcytosine (5mC), which has been extensively shown to play important roles in regulation of neural development, maturation, and plasticity (Gallegos et al., 2018). In addition to 5mC, DNA methylation at N6-deoxyadenosine (N6-mA) is the most prevalent DNA modification in prokaryotes (Alderman and Xiao, 2019; Heyn and Esteller, 2015). With the help of advancement of various deep sequencing techniques, recent studies started to discover N6-mA in eukaryotes, including human and rodents (Li et al., 2019; Li et al., 2020; Wu et al., 2016; Xie et al., 2018; Yao et al., 2017; Zhou et al., 2016). In several studies (Xiao et al., 2018; Xie et al., 2018; Zhou et al., 2016), ALKBH1 was identified to be the demethylase that erased DNA methylation at N6-deoxyadenosine. Importantly, in a latest co-crystal structure study (Zhang et al., 2020) mammalian ALKBH1 was revealed to serve as an N6-mA demethylase of unpairing DNA. In contrast to demethylase, the identities of the N6-mA methyltransferases in mammalian cells remain inconclusive. Either METTL4 or N6AMT1 has been suggested to be the putative DNA N6-mA methyltransferase (Kweon et al., 2019; Xiao et al., 2018).

Functionally, in mouse embryonic stem cells N6-mA has been shown to be enriched at LINE-1 transposon and function to silence gene transcription (Wu et al., 2016). In this study, ALKBH1 was identified to be the DNA N6-mA demethylase. In human mesenchymal stem cells (MSCs), decreased level of DNA N6-mA regulated by ALKBH1 has also been shown to be necessary for osteogenic differentiation of MSCs via promoting ATF4 transcription (Zhou et al., 2016). One recent study demonstrated that the level of N6-mA was drastically higher in human glioblastoma stem cells and primary human tumor samples (Xie et al., 2018), suggesting a functional role of DNA N6-mA in regulation of tumorigenesis. Indeed, silencing the demethylase ALKBH1 inhibited the proliferation of glioblastoma stem cells, whereas N6AMT1 did not seem to act as the methyltransferase in glioblastoma cells. Interestingly, N6-mA DIP sequencing (DIP-seq, DNA immunoprecipitation with anti-N6-mA antibody followed by next generation sequencing) revealed that many N6-mA peaks were enriched in genes regulating neurogenesis and neural development, suggesting that N6-mA function to repress these neurodevelopmental genes. This finding implied that DNA N6-mA might play important roles in neurons. Indeed, in the mouse brain the overall level of DNA N6-mA in the prefrontal cortex was found to be significantly elevated under stress condition (Yao et al., 2017). However, detailed genome wide analysis of N6-mA enrichment found that there was also region-specific loss of N6-mA. Consistent with the finding from glioblastoma cells (Xie et al., 2018), transcriptomic profiling upon stress demonstrated that upregulated genes associated with region-specific reduced level of N6-mA were related to neuronal development, such as neurogenesis and axon development. Furthermore, a new study (Li et al., 2019) discovered that upon fear extinction training, the level of DNA N6-mA in cortical neurons was significantly increased. The accumulation of N6-mA in response to the fear extinction training was due to the elevated expression of the methyltransferase N6AMT1. In contrast to many other studies, in which DNA N6-mA acts to silence gene transcription, this study showed that N6-mA functioned to promote the transcription of BDNF.

Axonal injury and regeneration are well known to involve neuronal stress responses and changed neural activity. Here we examined the roles of ALKBH1 in regulation of peripheral nerve regeneration. We found that ALKBH1 functioned as a demethylase in mouse adult sensory neurons and its mRNA level was significantly increased upon peripheral nerve injury. Knocking down ALKBH1 resulted in impaired sensory axon regeneration *in vitro* and *in vivo*, which could be rescued by simultaneously knocking down N6AMT1. Moreover, we observed that deleting ALKBH1 resulted in down regulation of several neurodevelopmental related genes, such as Id1 and neuritin, proteins known to regulate axon growth and regeneration (Gao et al., 2016; Huang et al., 2020; Huang et al., 2019; Lasorella et al., 2005; Sharma et al., 2015; Zhao et al., 2018). Collectively, our study not only discovered a new physiological function of ALKBH1 in neurons, but also illustrated a novel epigenetic regulatory mechanism underlying mammalian axon regeneration.

## Materials and methods

### Animals

All experiments using live rodents were performed in accordance with the animal protocol approved by the Institutional Animal Care and Use Committee of the Johns Hopkins University. The female CF-1 mice (6-week-old, weighing from 30 to 35 g) were ordered from Charles River Laboratories and housed in the institutional animal maintenance facility.

### Primary culture of sensory DRG neurons

Primary culture of DRG neurons from adult mice was performed according to the procedures described in previous studies (Hur et al., 2011a; Hur et al., 2011b; Saijilafu et al., 2013; Wang et al., 2018). Specifically, the DRGs from 6- to 8-week-old female CF-1 mice were dissected out, and then digested to isolating cells with type-I collagenase (1 mg/mL, Thermo Fisher Scientific) and dispase II (5 mg/mL, Thermo Fisher Scientific) at 37°C for 60 min. Cells were then washed 3 times with MEM containing 10% fetal bovine serum and 1X penicillin/streptomycin (Thermo Fisher Scientific) to terminate the enzymatic digestion. The digested DRGs were then dissociated into single cells with pipette trituration in the same medium and filtered through a 70-µm cell strainer. After centrifugation at 1000 rpm for 7 minutes, the cell pellets were then resuspended for electroporation (see below). After electroporation, the cells were immediately transferred into pre-warmed MEM medium and seeded onto glass coverslips coated with a mixture of 100 μg/mL poly-D-lysine (Sigma-Aldrich) and 10 μg/mL laminin (Thermo Fisher Scientific). After the neurons were fully settled and attached to the coverslips (about 6 hours after seeding), the medium was gently replaced with the culture medium, which was composed of MEM containing 5% fetal bovine serum, 1X GlutaMAX-I (Thermo Fisher Scientific), 1X penicillin/streptomycin and anti-mitotic reagents (20 μM 5-fluoro-2-deoxyuridine and 20 µM uridine, both from Sigma-Aldrich). The medium replacement removed the electroporation buffer and dead cells.

### *In vitro* electroporation of adult sensory neurons

The dissociated sensory neuron pellet was resuspended with 98 µL electroporation buffer (Mouse neuron nucleofector kit, Lonza) mixed with RNA oligos (0.2 nmol per transfection) and/or GFP plasmid (5-10 µg). The mixed solution was then transferred into a 2.0-mm electroporation cuvette, and the electroporation was performed using the Nucleofector II (Lonza) programmed G-013 for mouse neurons.

### Sensory neuron resuspension and replating

For RNAi-mediated loss-of-function experiments, we used a replating procedure used in one of our previous published studies (Saijilafu et al., 2013). Briefly, after *in vitro* electroporation of siRNAs the neurons were seeded directly onto poly-D-lysine/laminin coated plastic culture plate at much higher density. Three days later the cultured neurons were re-suspended gently with 1 ml pipette and replated onto glass coverslips coated with a mixture of 100 μg/ml poly-D-lysine (Sigma-Aldrich) and 10 μg/ml laminin (Thermo Fisher Scientific). Three days after transfection of siRNAs, the protein of interest should be significantly reduced in neurons. Therefore, the resuspension and replating approach would allow axons to regrow anew from neurons in the absence of the targeted protein. After 24 hours in culture, the replated neurons were fixed with 4% paraformaldehyde (PFA) for immunocytochemistry study.

### Immunocytochemistry of sensory neurons

Cultured adult sensory neurons were fixed with 4% (w/v) paraformaldehyde (PFA) at room temperature for 15 min and then washed with 1X phosphate-buffered saline (PBS). Fixed cells were then incubated with the blocking buffer, which was composed of 10% goat serum, 0.1% Triton X-100 in 1X PBS, for 60 min in dark at room temperature. Incubation with the primary antibodies diluted in blocking buffer were performed overnight at 4°C. After washing with PBS for three times, the secondary antibody incubation was performed in dark at room temperature for 60 min. After washing with 1X PBS 3 times and distilled water for one time, the coverslips were mounted onto glass slides with Fluoroshield histology mounting (Sigma-Aldrich).

### Imaging and analysis of *in vitro* regenerative axon growth of sensory neurons

Fluorescent image tiles of the whole coverslip were captured with a CCD camera connected to an inverted fluorescent microscope controlled by AxioVision (Zeiss) using the MosaiX module. The tiles were then stitched into a single image. The image was exported in grey scale Tiff format and modified with ImageJ software in accordance with the preference of Neuron Image Analyzer (Kim et al., 2015), which is a program running on the MATLAB platform to identify cell somas and trace the longest process from each soma. The code was modified slightly to suit our tracing purpose. Percentage of Neurons with Axon was calculated as the ratio of Traced Neurons vs. Identified Neurons.

### *In vivo* electroporation of adult sensory neurons

The *in vivo* electroporation of adult mouse DRGs was performed as previously described (Li et al., 2018; Saijilafu et al., 2011). In brief, after a mouse was stably anesthetized, a L3-5 dorsolateral laminectomy was performed on the left side to expose the left L4-5 DRGs. 1-1.5 µL solution containing desired plasmids (2.0 µg/µL per transfection) and/or RNA oligos (0.1 nmol per transfection) were injected into each DRG with a glass micropipette pulled by a glass capillary puller (NARISHEGE) connected to the Picospritzer III system (30-psi pressure, 8-ms duration, Parker Instrumentation). *In vivo* electroporation was performed immediately after the injection by applying five square electric pulses (35 V, 15-ms duration, 950-ms interval) using a platinum tweezertrode (BTX Harvard Apparatus) powered by the ECM 830 square wave electroporation system (BTX Harvard Apparatus). The cut wound was then closed. Two days after the mouse was recovered from the surgery, a small cut was made on the left thigh to expose the sciatic nerve, which was then crushed with Dumont #5 forceps. A small knot was made on the crush site with 10-0 nylon epineural sutures as a mark. After 2 or 3 days, the mouse was perfused transcardially with 1X PBS followed by 4% PFA. The sciatic nerve segment including the crush site was then dissected out and post-fixed in 4% PFA overnight at 4°C. The fixed nerve segment was then mounted onto a glass slide and covered with a coverslip and flattened with pressure before imaging. The siRNAs targeting mouse ALKBH1, N6AMT1 and non-targeting siRNA control were purchased from Dharmacon. All target sequences of siRNAs used are listed in Table S1.

### Imaging and analysis of *in vivo* sensory axon regeneration

For quantification of *in vivo* axon regeneration, the images of whole-mount sciatic nerve segments were first captured with a CCD camera connected to an inverted fluorescence microscope controlled by the AxioVision software (Zeiss) using the MosaiX module by tiles. The tiles were stitched into a single image with AxioVision software (Zeiss). Each axon with identifiable growth tip in the nerve segment was manually traced from the crush site to the distal axonal tip with the same software to determine the length. The mean of all traced axons in one nerve image was used as the average axon length of this nerve. All the traced axons lengths within one experimental group were used to calculate the cumulative distribution. Nerve images with less than 15 tracible axons were excluded from data analysis. Experimenters quantifying the axon lengths were blinded to conducted the experimental conditions. The *in vivo* plasmids and siRNA transfection rates were quantified in a previous study (Li et al., 2018).

### Quantitative real-time PCR

Total RNA was isolated from cells or tissues according to the protocols in RNeasy Plus mini kit (QIAGEN), and then reversely transcribed to first strand cDNA using SuperScript™ III First-Strand Synthesis SuperMix for qRT-PCR (Invitrogen). For real-time qPCR analysis, 10 ng first strand cDNA was amplified with gene-specific primers and PowerUp™ SYBR™ Green Master Mix (Applied Biosystems) using the StepOnePlus™ Real-Time PCR System (Applied Biosystems). All experiments were repeated for three times. Relative expression levels of mRNAs were calculated using the 2-ΔΔCt method and normalized to control group respectively. Gapdh was used as the endogenous control for all mRNAs. All the primers used are listed in Table S2.

### DNA dot blot

Genomic DNA from CAD cells or sensory neurons was isolated according to the protocol of the PureLink Genomic DNA kit (Invitrogen). For dot blotting, the genomic DNA was denatured at 95 °C for 10 min in 0.4 mol/L NaOH and 10 mmol/L EDTA buffer. Samples were spotted on the membrane (Zeta-Probe, Bio-Rad, Hercules, CA, USA) using a Dot Blot micro-filtration apparatus (Bio-Rad) and cross-linked by UV Stratalinker 1800 (Stratagene) two times. Membrane was then blocked in blocking buffer (5% milk in TBST) for 1 h at room temperature and incubated with N6-mA specific antibody (Synaptic Systems, 1:2000) overnight at 4 °C. After 3 times washing with TBST, the membranes were incubated with HRP-linked horse anti-rabbit IgG (Cell Signaling Technology). The antibody-antigen complexes were visualized with Kwik Quant Imager (Kindle Biosciences). To ensure an equal amount of DNA was spotted, the same membrane was stained with 0.02% methylene blue in 0.3 mol/L sodium acetate (pH 5.2). RNAase treatments were used to ensure the complete removal of RNA contamination. To quantify the DNA dot blot results, we used Image J software to measure the total density of the N6-mA signal normalized to the corresponding meth-blue signal.

### Statistical analysis

Data were analyzed with GraphPad Prism 8 and presented as mean ± SEM unless specifically stated. The significance level was set as p < 0.05. For comparisons between two groups, regular two-tailed Student’s t test was used to determine the statistical significance. If the data were normalized to the control group as relative levels, one sample t test with 1 as the hypothetical value was used. For comparisons among three or more groups, one-way ANOVA followed by Turkey’s multiple comparison test was used to determine the statistical significance. All statistical details of experiments, including the statistical tests used, exact value of n, definition of n, can be found in figure legends.

## Results

### ALKBH1 regulates N6-mA DNA methylation in mammalian sensory neurons

We first measured the levels of N6-mA using a N6-mA specific antibody validated in several previously published studies (Xie et al., 2018; Yao et al., 2017; Zhou et al., 2016). Genomic DNAs isolated from Hela cells, CAD cell which is a mouse neuronal cell line, and primary sensory neurons from the dorsal root ganglion (DRG) were used. The dot blot results showed that anti-N6-mA antibody could detect DNA N6-mA, showing clear signals in a concentration-dependent manner (Figure 1A). To examine if ALKBH1 acted as a N6-mA demethylase in neurons, we first used siRNAs against ALKBH1 in differentiated neuronal CAD cells and then measured the levels of N6-mA. The dot blot results showed that compared with the cells transfected with non-targeting siRNAs (siNon-Target), knocking down ALKBH1 in CAD cells with siRNAs (siALKBH1) resulted in significantly higher level of N6-mA. Moreover, knocking down both ALKBH1 and the DNA N6-mA methyltransferase N6AMT1 (siN6AMT1) reduced the level of N6-mA back to the control condition (Figure 1B, C). These results provided strong evidence that ALKBH1 functioned as a N6-mA demethylase in CAD cells, removing DNA N6-mA group which was methylated by N6AMT1. Interestingly, knocking down N6AMT1 alone did not significantly decrease the level of N6-mA (Figure 1B, C), suggesting that the endogenous level of DNA N6-mA in neuronal cells is already low in the control condition. Indeed, with similar amount of isolated DNA, the N6-mA level in Hela cells was much higher than that in CAD cells (Figure 1A, D). To confirm that the detected N6-mA dot signals were indeed from genomic DNAs rather than RNA N6 modifications, we treated the isolated samples with DNase A before the dot blot experiments. The results showed that DNase A almost completely eliminate the meth-blue and dot blot signals detected with the specific N6-mA antibody (Figure 1D).

**Figure 1.**
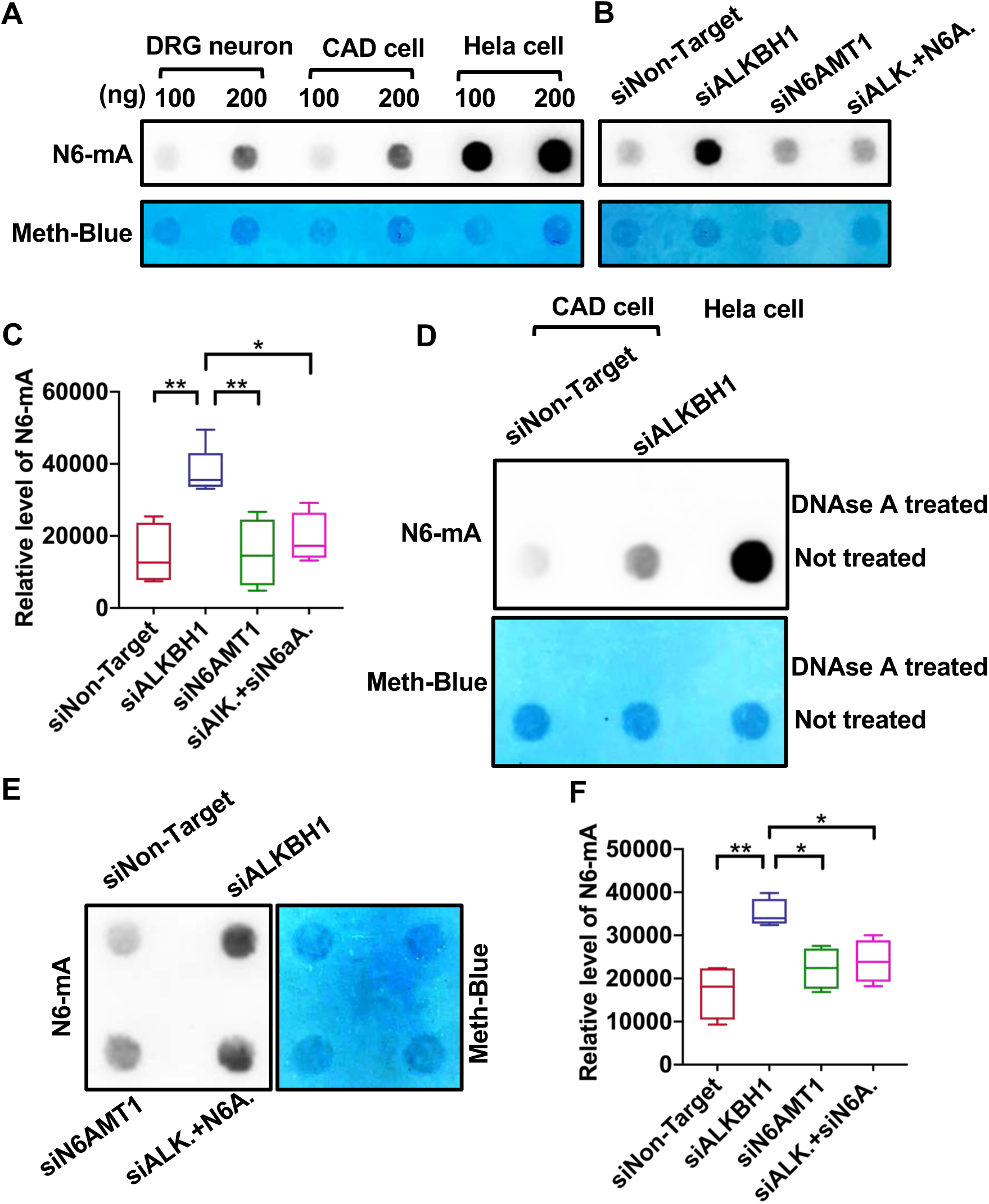
ALKBH1 acted as a DNA N6-deoxyadenosine (N6-mA) demethylase in mouse sensory neurons. (A) Representative images of dot blot assay showing that the specific antibody against DNA N6-mA can detect the levels of N6-mA in mouse DRG neurons, the differentiated neuronal cell line CAD cells, and the Hela cells, in a dose-dependent manner. (B) Representative dot blot images showing that knocking down ALKBH1 in CAD cells led to increased level of N6-mA, whereas knocking down N6AMT1 alone had little effect. However, when ALKBH1 and N6AMT1 were knocked down together, the level of N6-mA was reversed back to the control condition. (C) Quantification of (B) (one-way ANOVA followed by Tukey’s multiple comparisons test, *P* = 0.0013, n = 4-5 independent experiments). (D) Representative dot blot images showing that treating the isolated DNA samples with DNase resulted in total loss of sample signals and associated N6-mA signals in either CAD or Hela cells. (E) Representative dot blot images showing that knocking down ALKBH1 in adult mouse DRG neurons led to increased level of N6-mA, whereas knocking down N6AMT1 alone had little effect. However, when ALKBH1 and N6AMT1 were knocked down together, the level of N6-mA was reversed back to the control condition. (F) Quantification of (B) (one-way ANOVA followed by Tukey’s multiple comparisons test, *P* = 0.002, n = 4 independent experiments). Data are represented as mean ± SEM. *P* < 0.05, \*\**P* < 0.01, \*\*\**P* < 0.001, \*\*\*\**P* < 0.0001, compared to control if not designated.

We next performed the same sets of experiments using cultured adult mouse primary sensory neurons. The results showed that knocking down ALKBH1 also led to significantly increased level of N6-mA, which was significantly reduced by co-knocking down N6AMT1 (Figure 1E, F). These results indicated clearly that ALKBH1 acted as a DNA N6-mA demethylase in adult mouse sensory neurons.

### ALKBH1 is necessary for regenerative sensory axon growth *in vitro*

To investigate if ALKBH1 functioned in sensory neurons to regulate regenerative axon growth, we used our previously established culture-replate model (Saijilafu et al., 2013). Such approach allowed us to evaluate regenerative axon regrowth from the neuronal soma after the genes of interests have been knocked down. Specifically, adult mouse sensory neurons were first transfected with either siNon-Target, siALKBH1, siN6AMT1, or siALKBH1+N6AMT1 via electroporation and culture for 3 days. The real time PCR results showed that siALKBH1 or siN6AMT1 transfection resulted in significantly reduced level of ALKBH1 or N6AMT1 in sensory neurons after 3 days in culture (Figure 2A, B). The neurons were then re-suspended and replated for another 24 hours before fixation (Figure 2C). Quantification of the average lengths of axons 24 hours after neuronal replating demonstrated that knocking down ALKBH1 significantly impaired regenerative axon growth (Figure 2C, D). Although knocking down N6AMT1 alone had little effect, it significantly rescued axon growth defect induced by down regulation of ALKBH1 (Figure 2C, D). When the percentages of neurons with axons were quantified, the results showed that down regulation of ALKBH1 and/or N6AMT1 had no effects, suggesting that they do not regulate axon initiation from the neuronal soma (Figure 2E). Together, these results demonstrated that ALKBH1 and its regulated DNA N6-mA demethylation were necessary for sensory axon regeneration *in vitro*.

**Figure 2.**
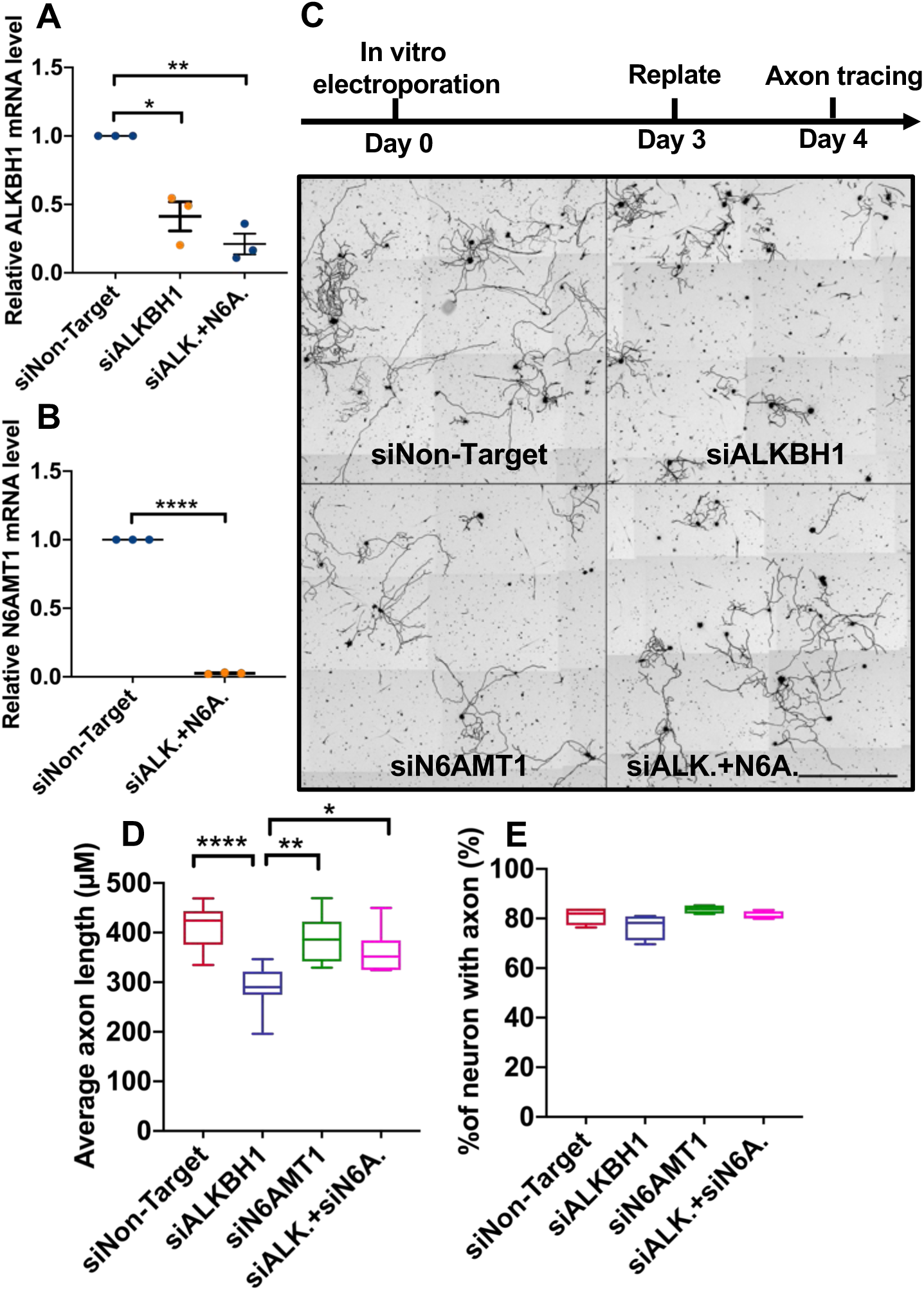
ALKBH1 and N6AMT1 regulated regenerative axon growth of sensory neurons *in vitro*. (A) Real-time PCR analysis showing significantly reduced mRNA level of ALKBH1 3 days after electroporation of siRNAs against ALKBH1 (siALKBH1) or together with siRNAs against N6AMT1 (siALK.+siN6A.). One sample t test, *P* = 0.0315 and 0.0092 for siALKBH1 and siALK.+siN6A., respectively. n = 3 independent experiments. (B) Real-time PCR analysis showing significantly reduced mRNA level of N6AMT1 3 days after electroporation of siRNAs against ALKBH1 and N6AMT1. One-sample t test, *P* < 0.0001, n = 3 independent experiments. (C) Top: time line of the culture and replate experiments. Bottom: representative images of cultured sensory neurons 24 hours post-replating after knocking down ALKBH1, N6AMT1, or together with siRNAs (siALKBH1, siN6AMT1, or siALK.+siN6A.). Scale bar, 500 μm. (D) Quantification of average lengths of the longest neurites (one-way ANOVA followed by Tukey’s multiple comparisons test, *P* < 0.0001, n = 7-10 independent experiments). (E) Quantification of percentage of neurons with axons (one-way ANOVA followed by Tukey’s multiple comparisons test, *P* = 0.0827, n = 4 independent experiments). Data are represented as mean ± SEM. *P* < 0.05, \*\**P* < 0.01, \*\*\**P* < 0.001, \*\*\*\**P* < 0.0001, compared to control if not designated.

### ALKBH1 is necessary for fast sensory axon regeneration *in vivo*

We first examined the expression levels of ALKBH1 in sensory neurons after peripheral nerve injury. The result showed that the mRNA level of ALKBH1 was significantly elevated 1 day after peripheral nerve injury and returned back to the control level after 3 days (Figure 3A). To evaluate if ALKBH1 played a role in regulation of sensory axon regeneration *in vivo*, we performed *in vivo* electroporation with either siNon-Target or siALKBH1 into adult sensory neurons and performed sciatic nerve crush injury 2 or 3 days after the electroporation. Real-time PCR data showed that 3 days after *in vivo* electroporation of siALKBH1, its mRNA level in DRGs was significantly reduced (Figure 3B) compared with that in control DRGs (transfected with siNon-Target). An early study (Smith and Skene, 1997) demonstrated that, the sensory neurons switched into fast axon regeneration mode about 2 days after axotomy. In consistent, our recent time-course analysis of sensory axon regeneration (Gao et al., 2020) showed that the rate of axon regeneration reached the fastest rate about 3 days after the sciatic nerve injury. Therefore, we examined sensory axon regeneration at 2, 3, or 4 days after the nerve crush. The results showed that 2 days after the nerve crush when axon regeneration rate was still low, knocking down ALKBH1 did not have significant effect on axon regeneration (Figure 3C-E). Similarly, knocking down N6AMT1 also had no effect on sensory axon regeneration 2 days post-nerve injury (Figure 3F-H). In contrast, when sensory axon regeneration was examined at 3 days after the nerve crush, knocking down ALKBH1 led to significantly reduced sensory axons’ spontaneous regeneration (Figure 4A-C), indicating that ALKBH1 was necessary for fast sensory axon regeneration *in vivo*. Moreover, knocking down ALKBH1 still impaired sensory axon regeneration 4 days after the nerve injury (Figure 4D-F), suggesting that the inhibitory effect is sustained.

**Figure 3.**
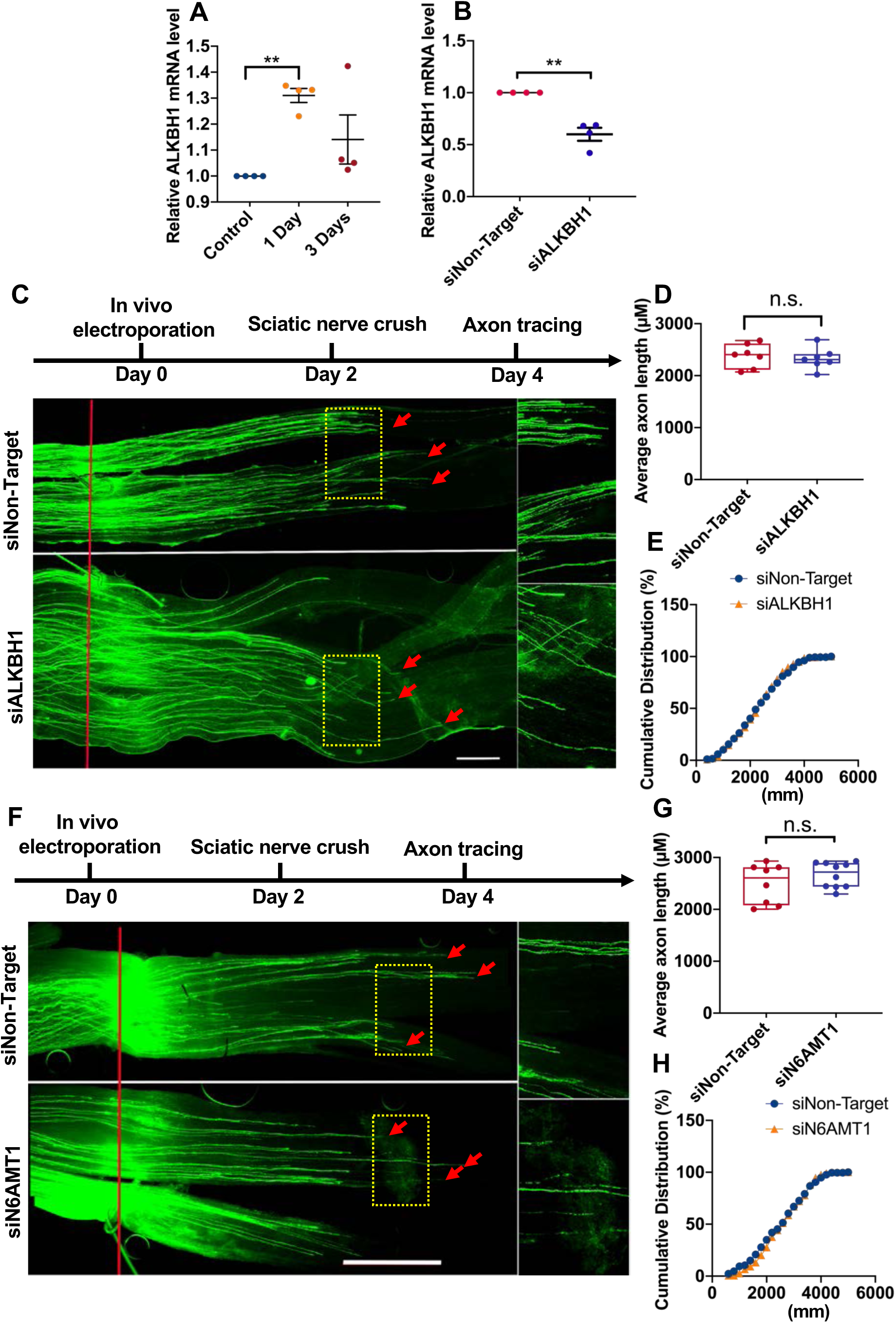
Down-regulation of ALKBH1 or N6AMT1 in sensory neurons had no effect on initial slow sensory axon regeneration *in vivo*. (A) Real-time PCR analysis showing significantly elevated mRNA level of ALKBH1 in sensory neurons 1-day post-nerve injury (one sample t test, *P* = 0.0014, n = 4 independent experiments). (B) Real-time PCR analysis showing significantly reduced mRNA level of ALKBH1 *in vivo* after 3 days after electroporation (one sample t test, *P* = 0.0076, n = 4 independent experiments). (C) Top: timeline of the experiment. Bottom: representative images showing that knocking down ALKBH1 had no effect on slower sensory axon regeneration 2 days post-nerve injury. The right column shows enlarged images of areas in the dashed yellow boxes. The red line indicates the crush sites and the read arrows label regenerating axon tips. Scale bar, 1 mm for the left panel and 0.5 mm for the right panel. (D) Quantification of average length of regenerating sensory axons showing no significant difference between the control group and the ALKBH1 knockdown group 2 days post-nerve injury (unpaired student t test, *P* = 0.6387, n = 7 mice in each condition). (E) Cumulative distribution curves showing similar axon regeneration between the control group and the ALKBH1 knockdown group 2 days post-nerve injury. (F) Top: timeline of the experiment. Bottom: representative images showing that knocking down N6AMT1 had no effect on slower sensory axon regeneration 2 days post-nerve injury. The right column shows enlarged images of areas in the dashed yellow boxes. The red line indicates the crush sites and the read arrows label regenerating axon tips. Scale bar, 1 mm for the left panel and 0.5 mm for the right panel. (G) Quantification of average length of regenerating sensory axons showing no significant difference between the control group and the N6AMT1 knockdown group 2 days post-nerve injury (unpaired student t test, *P* = 0.2715, n = 7 and 10 mice in control and siN6AMT1 group, respectively). (H) Cumulative distribution curves showing similar axon regeneration between the control group and the N6AMT1 knockdown group 2 days post-nerve injury. Data are represented as mean ± SEM. *P* < 0.05, \*\**P* < 0.01, \*\*\**P* < 0.001, \*\*\*\**P* < 0.0001, compared to control if not designated.

**Figure 4.**
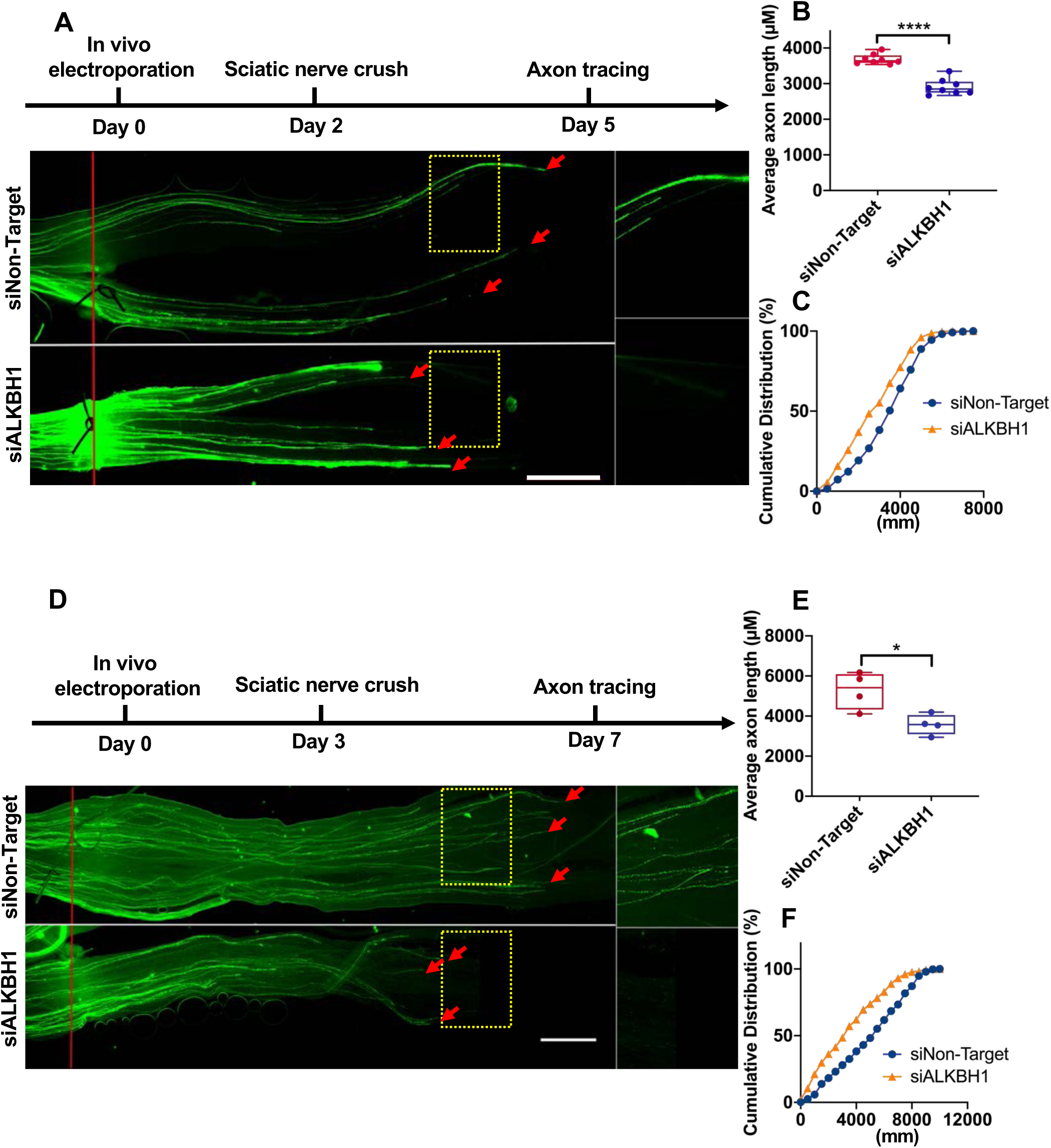
Down-regulation of ALKBH1 in sensory neurons impaired fast sensory axon regeneration *in vivo*. (A) Top: timeline of the experiment. Bottom: representative images showing that knocking down ALKBH1 significantly impaired fast sensory axon regeneration 3 days post-nerve injury. The right column shows enlarged images of areas in the dashed yellow boxes. The red line indicates the crush sites and the read arrows label regenerating axon tips. Scale bar, 1 mm for the left panel and 0.5 mm for the right panel. (B) Quantification of average length of regenerating axons showing that knocking down ALKBH1 significantly impaired sensory axon regeneration 3 days post-nerve injury (unpaired student t test, *P* < 0.0001, n = 8 mice in each condition). (C) Cumulative distribution curves showing that ALKBH1 knockdown resulted in significant reduced sensory axon regeneration 3 days post-nerve injury. (D) Top: timeline of the experiment. Bottom: representative images showing that knocking down ALKBH1 significantly impaired fast sensory axon regeneration 4 days post-nerve injury. The right column shows enlarged images of areas in the dashed yellow boxes. The red line indicates the crush sites and the read arrows label regenerating axon tips. Scale bar, 1 mm for the left panel and 0.5 mm for the right panel. (E) Quantification of average length of regenerating axons showing that knocking down ALKBH1 significantly impaired sensory axon regeneration 4 days post-nerve injury (unpaired student t test, *P* = 0.018, n = 4 mice in each condition). (F) Cumulative distribution curves showing that ALKBH1 knockdown resulted in significant reduced sensory axon regeneration 4 days post-nerve injury. Data are represented as mean ± SEM. *P* < 0.05, \*\**P* < 0.01, \*\*\**P* < 0.001, \*\*\*\**P* < 0.0001, compared to control if not designated.

We next examined how down regulation of N6AMT1 affected impaired sensory axon regeneration induced by ALKBH1 knockdown. First, we found that knocking down N6AMT1 still had little effect on sensory axon regeneration 3 days after the nerve crush (Figure 5), indicating that N6AMT1 is not necessary for spontaneous PNS axon regeneration. However, when ALKBH1 and N6AMT1 were knocked down together and sensory axon regeneration was examined 3 days after the nerve injury, the results showed that N6AMT1 knockdown significantly rescued sensory axon regeneration impaired by ALKBH1 deletion (Figure 5).

**Figure 5.**
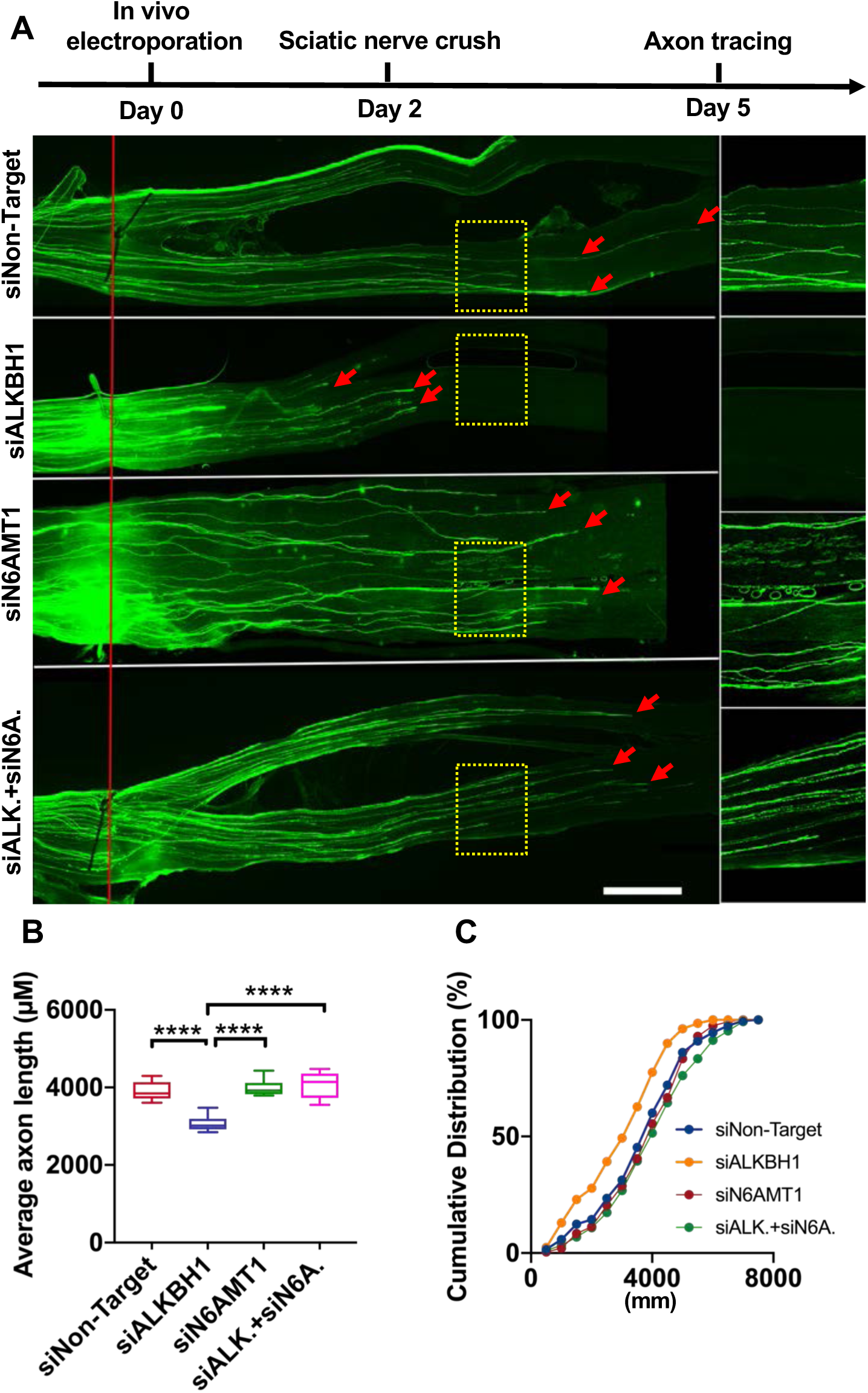
Down-regulation of N6AMT1 in sensory neurons reversed sensory axon regeneration impaired by knocking down ALKBH1. (A) Top: timeline of the experiment. Bottom: representative images showing that knocking down N6AMT1 reversed sensory axon regeneration impaired by down regulation of ALKBH1 3 days post-nerve injury. The right column shows enlarged images of areas in the dashed yellow boxes. The red line indicates the crush sites and the read arrows label regenerating axon tips. Scale bar, 1 mm for the left panel and 0.5 mm for the right panel. (B) Quantification of average length of regenerating axons under conditions shown in (A). One-way ANOVA followed by Tukey’s multiple comparisons test, *P* < 0.0001, n = 7-9 mice in each condition. (C) Cumulative distribution curves showing that knocking down N6AMT1 rescued axon regeneration impaired by ALKBH1 knockdown. Data are represented as mean ± SEM. *P* < 0.05, \*\**P* < 0.01, \*\*\**P* < 0.001, \*\*\*\**P* < 0.0001, compared to control if not designated.

In summary, these results not only provided evidence that ALKBH1 was necessary for fast sensory axon regeneration *in vivo*, but also suggested that increased level of DNA N6-mA in sensory neurons function to repress sensory axon regeneration.

### Down regulation of ALKBH1 resulted in changed expression of neurodevelopmental related genes

Several previous studies (Xie et al., 2018; Yao et al., 2017) in neurons or glia cells have shown that ALKBH1 and N6-mA functioned to repress the expression of many neurodevelopmental regulatory genes. Based on these studies, we first performed gene ontology (GO) analysis on a public RNA-seq dataset (GSE117632) acquired from patient-derived human glioblastoma cells with or without Alkbh1-knockdown (Xie et al., 2018). When running GO analysis on the differentially expressed genes (RPKM>1) with two-fold or greater downregulation upon Alkbh1-knockdown, the biological process termed regulation of neuron differentiation (GO: 0045664) enriched. Among these 65 genes, we selected 18 gene- of-interest for real-time qPCR analysis in mouse DRG neurons.

The results showed that after knocking down ALKBH1, among 18 genes the mRNA levels of 5 genes were significantly reduced compared to that of control neurons (siNon-Target), including *Efna1, Id1, Nrn1, ATG9B*, and *C1QL4* (Figure 6A). None of the examined genes was statistically significantly up regulated upon ALKBH1 knockdown, consistent with the idea that DNA N6-mA mainly functions to condense chromatin and repress gene expression. Interestingly, several of these 5 genes have been shown to regulate axon growth or regeneration. For instance, Nrn1 that encodes the protein neuritin is well known to promote axon growth and optic nerve regeneration (Gao et al., 2016; Huang et al., 2020; Sharma et al., 2015; Zhao et al., 2018). In addition, the transcription factor Id1 and its homolog Id2 have been shown to be involved in regulation of axon growth and regeneration (Huang et al., 2019; Lasorella et al., 2005). When N6AMT1 was knocked down together with ALKBH1, based on one-way ANOVA analysis, the levels of 4 genes were significantly affected, in which the levels of C1QL4 and Id1 were significantly reversed (Figure 6B-E). These results suggested that ALKBH1 might support sensory axon regeneration by reducing neuronal DNA N6-mA level and the subsequent up regulation of these neurodevelopmental genes.

**Figure 6.**
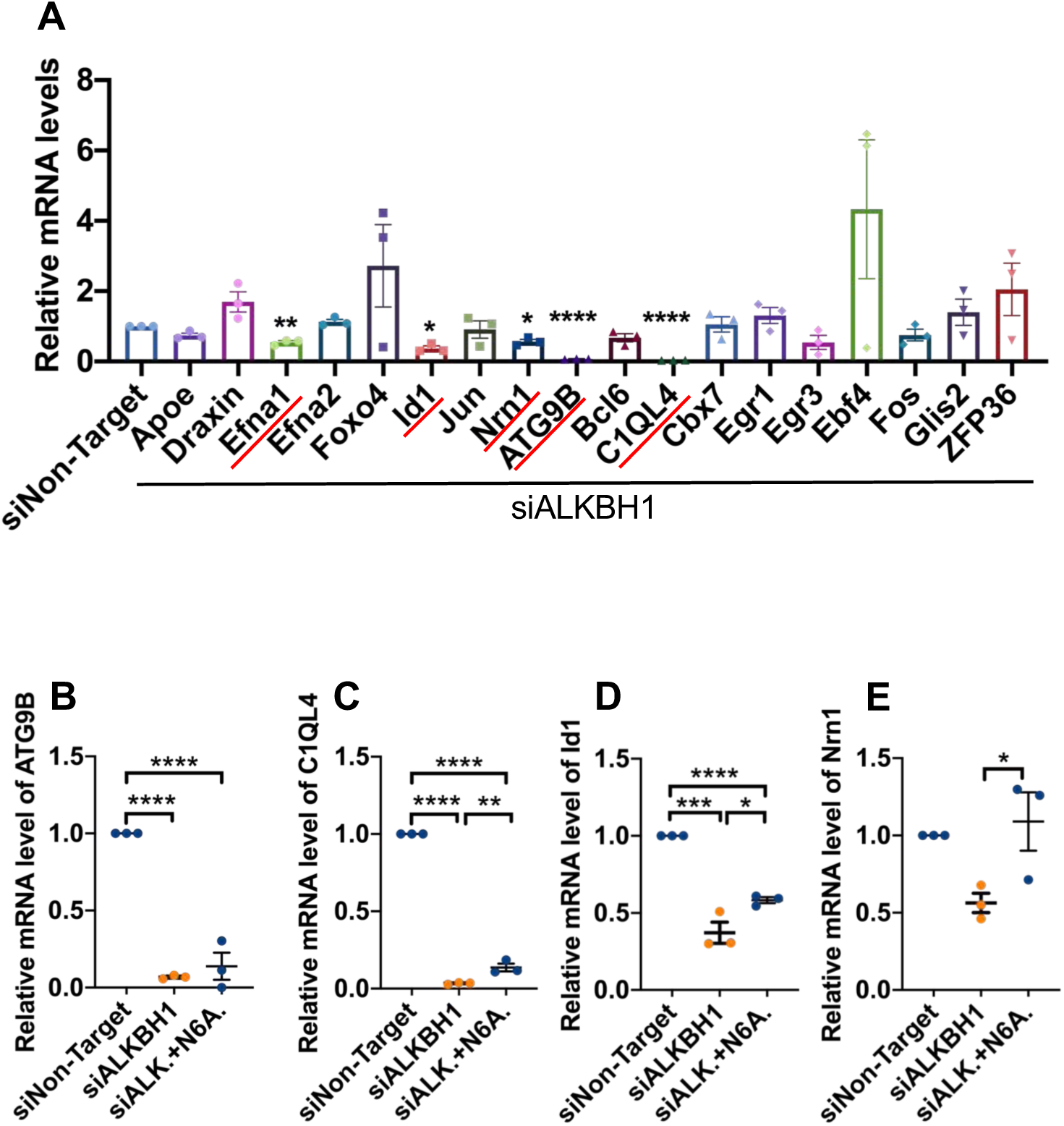
Knocking down ALKBH1 in sensory neurons led to down-regulation of neurodevelopmental related genes. (A) Real-time PCR analyses of changed transcription of 18 genes in sensory neurons after knocking down ALKBH1. The 5 significantly down regulated genes were underlined in red. One sample t test, *P* = 0.0096, 0.0118, and 0.0202 for *Efna1, Id1, and Nrn1*, respectively. *P* < 0.0001 for *ATG9B* or *C1QL4*. n = 3 independent experiments. (B) Real-time PCR analysis of *ATG9B* mRNA levels after knocking down ALKBH1 or double knocking down ALKBH1 and N6AMT1. One-way ANOVA followed by Tukey’s multiple comparisons test, *P* < 0.0001, n = 3 independent experiments. (C) Real-time PCR analysis of *C1QL4* mRNA levels after knocking down ALKBH1 or double knocking down ALKBH1 and N6AMT1. One-way ANOVA followed by Tukey’s multiple comparisons test, *P* < 0.0001, n = 3 independent experiments. (D) Real-time PCR analysis of *Id1* mRNA levels after knocking down ALKBH1 or double knocking down ALKBH1 and N6AMT1. One-way ANOVA followed by Tukey’s multiple comparisons test, *P* = 0.0001, n = 3 independent experiments. (E) Real-time PCR analysis of *Nrn1* mRNA levels after knocking down ALKBH1 or double knocking down ALKBH1 and N6AMT1. One-way ANOVA followed by Tukey’s multiple comparisons test, *P* = 0.0368, n = 3 independent experiments. Data are represented as mean ± SEM. *P* < 0.05, \*\**P* < 0.01, \*\*\**P* < 0.001, \*\*\*\**P* < 0.0001, compared to control if not designated.

## Discussion

There is emerging evidence that the intrinsic ability of mammalian axon regeneration is epigenetically regulated (Qian and Zhou, 2020), including microRNAs (Jiang et al., 2015; Liu et al., 2013; Wang et al., 2018), histone modifications (Cho and Cavalli, 2014; Finelli et al., 2013; Liu et al., 2013; Puttagunta et al., 2014), and DNA methylation (Oh et al., 2018; Weng et al., 2017). Neurons in the mammalian PNS are well known to regenerate robustly after peripheral nerve injury, making them a perfect model system to investigate the molecular mechanisms by which axon regeneration is regulated. In this study we demonstrated that sensory axon regeneration was regulated by ALKBH1, recently identified to be the demethylase of DNA N6-mA (Li et al., 2020; Xiao et al., 2018; Zhang et al., 2020), providing an important physiological function of ALKBH1 and DNA N6-mA modification.

Although many recent studies revealed the important roles of DNA N6-mA modification in regulation of chromatin structure and gene expression in mammals (Li et al., 2019; Li et al., 2020; Wu et al., 2016; Xie et al., 2018; Yao et al., 2017; Zhou et al., 2016), there still are debating regarding its existence in mammals (Liu et al., 2017; Ratel et al., 2006; Schiffers et al., 2017). For instance, a recent study (Douvlataniotis et al., 2020) suggested that contamination of bacterial in cultured cells or mRNA in obtained DNA samples might artificially contribute to the detection of DNA N6-mA signal. In our study, we provided evidence that knocking down ALKBH1 in 2 cell lines and 1 primary neuron led to significantly increased levels of DNA N6-mA. Moreover, co-deletion of N6AMT1 significantly reduced the levels of DNA N6-mA in CAD cells and sensory neurons, providing confirming evidence regarding the presence of DNA N6-mA in neurons. When obtained DNA samples were treated with DNase I, both methyl-blue and N6-mA signals were completely lost, indicating that mRNA contamination was unlikely. Together, our data indicated strongly that DNA N6-mA modification could be detected in sensory neurons, which was regulated by the demethylase ALKBH1 and the methyltransferase N6AMT1.

In our study, we provided both *in vitro* and *in vivo* data that sensory axon regeneration was supported by ALKBH1, likely mediated by its role as the DNA N6-mA demethylase. In addition, co-deletion of the methyltransferase N6AMT1 completely rescued axon regeneration impaired by ALKBH1 down-regulation both *in vitro* and *in vivo*, further confirming the important function of DNA N6-mA in regulation of sensory axon regeneration. In two recent studies, DNA N6-mA has been shown to regulate mouse neuronal gene expression in response to environmental stress (Yao et al., 2017) or neural activity (Li et al., 2019). In particular, in the first study the dynamic change of N6-mA in the mouse prefrontal cortex neurons was found to be linked to environmental stress. Importantly, the study revealed that loss of N6-mA was strongly associated with up regulation of genes regulating neuronal development, such as neurogenesis and axon development. Indeed, in our study we also observed that knocking down ALKBH1 (increased level of N6-mA) in adult mouse sensory neurons resulted in down regulation of neurodevelopmental genes. Conversely, peripheral nerve crush resulted in up regulation of ALKBH1 (decreased level of N6-mA), and thus presumably up-regulation of neurodevelopmental genes. Interestingly, a recent study (Palmisano et al., 2019) suggested that sensory axon regeneration shared similar molecular pathways with that during developmental axon growth. Another study (Poplawski et al., 2020) also showed that CNS neurons undergoing successful axon regeneration re-express many developmental related genes. Therefore, our results provided a potential mechanism underlying such rejuvenation of gene expression during axon regeneration. Indeed, as aforementioned we found that ALKBH1 supported the expression of Id1 and neuritin, whereas N6AMT1 had opposite effects. In other words, our results indicated that in adult mouse sensory neurons DNA N6-mA modification acted to repress the expression of neurodevelopmental genes supporting axon growth. It will be of great importance that future experiments are performed to see if overexpression of these genes (e.g. neuritin, Id) can rescue axon regeneration impaired by knocking down ALKBH1.

It should be noted that in our study, knocking down the methyltransferase N6AMT1 did not change the level of N6-mA or axon regeneration. However, knocking down N6AMT1 could significantly rescue ALKBH1 knockdown-induced elevated level of N6-mA and reduced axon regeneration. One potential explanation is that the endogenous level of DNA N6-mA in sensory neurons is relatively low comparing to other cell types and ALKBH1 is the major regulator of such DNA modification. Thus, it will be also very interesting in the future to determine if the level of DNA N6-mA in mature CNS neurons is high, which might underlie their low intrinsic regeneration ability, and if overexpression of ALKBH1 can enhance CNS axon regeneration.

## Author contributions

Q. L., C. Q. and F-Q. Z. designed the experiments. Q. L. and C. Q. performed the experiments and analyzed data. H. F., T. L. and Y. H. helped analyzing the data. Q. L. and F-Q. Z. co-wrote the paper with input from all authors.

## Acknowledgements

The study was supported by grants (to F.Q.Z.) from NIH (R01NS085176, R01GM111514, R01EY027347, R01EY030883, R01EY031779), the Craig H. Neilsen Foundation (259450), and the BrightFocus Foundation (G2017037).

---

**Table S1.**
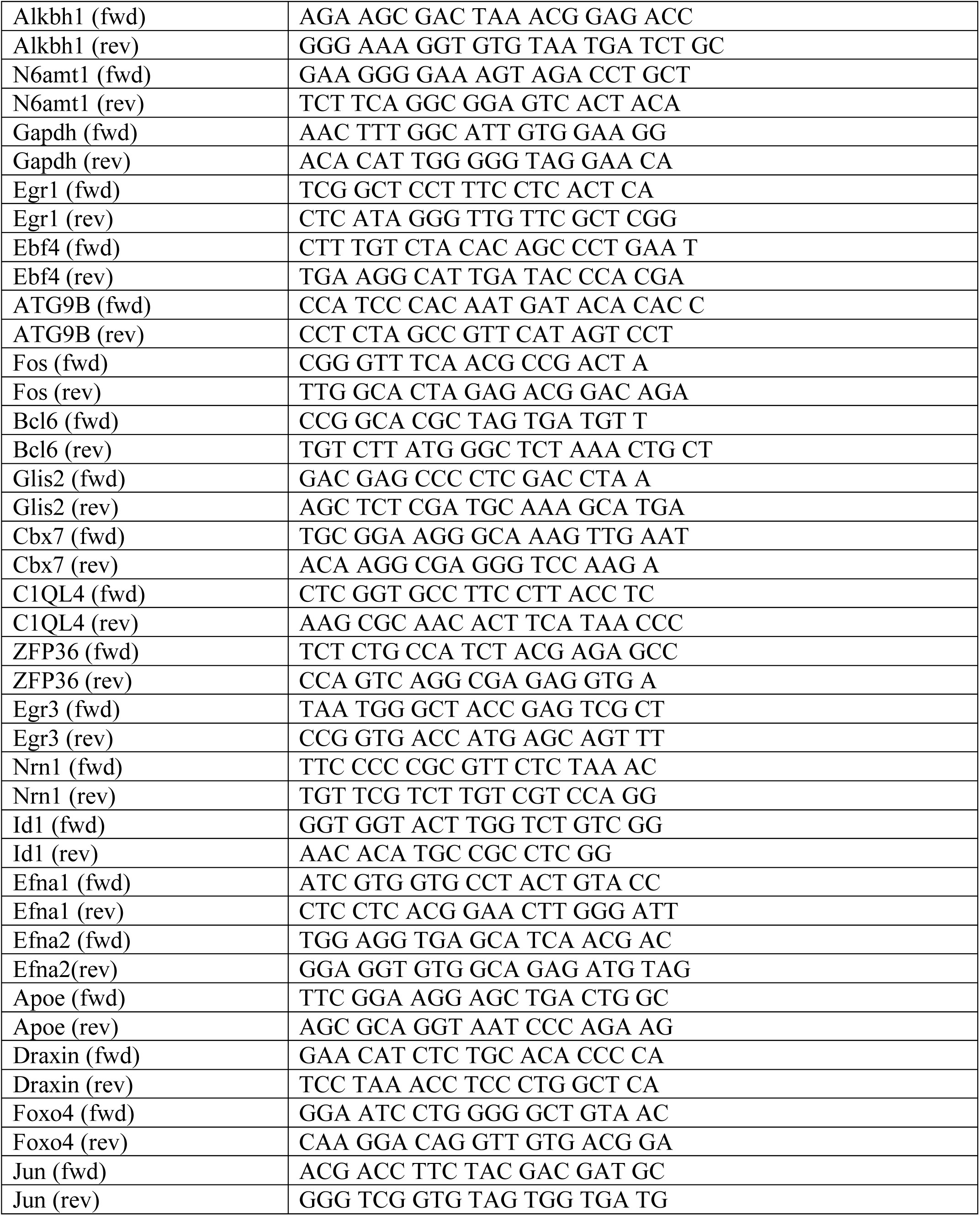
Primers for real-time PCR.

**Table S2.**
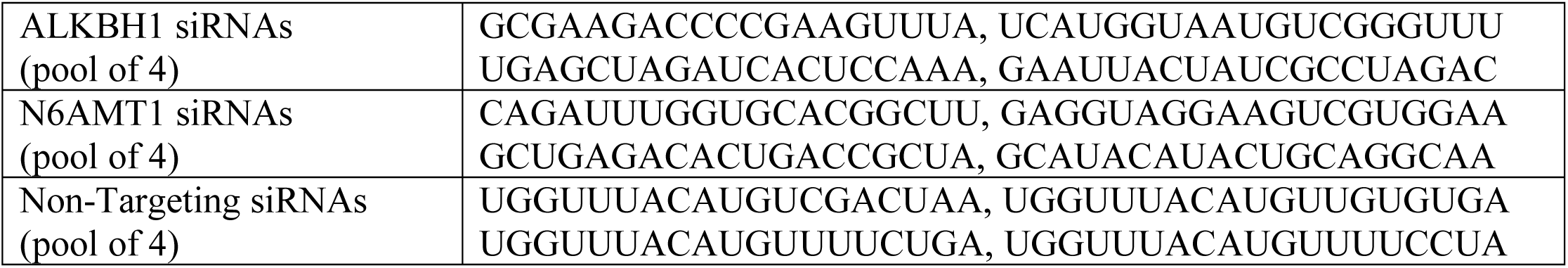
Target sequences of siRNAs.

